# Spatial distribution of lamin A determines nuclear stiffness and stress-mediated deformation

**DOI:** 10.1101/765263

**Authors:** Luv Kishore Srivastava, Zhaoping Ju, Ajinkya Ghagre, Allen J. Ehrlicher

**Affiliations:** Department of Bioengineering, McGill University, Montreal H3A 0E9; Department of Anatomy and Cell Biology, McGill University, Montreal H3A 0C7

## Abstract

The nucleus is the largest organelle and information center of the cell; while diverse cellular components have been identified as mechanotransduction elements, the deformation of the nucleus itself is emerging as a critical mechanosensory mechanism, suggesting that the nuclear stiffness is essential in determining responses to intracellular and extracellular stresses. The nuclear membrane protein, lamin A, is known to be a dominant component in nuclear stiffening; however, the quantitative relationship between lamin A expression and nuclear deformation is still unclear. Here we measure the nuclear moduli as a function of lamin A expression and applied stress, revealing a linear dependence of bulk modulus on lamin A expression. We also find that the bulk modulus is anisotropic, with the vertical axis of the nucleus being more compliant than the minor and major axis. To examine how lamin A influences nuclear mechanics at the sub-micron scale we correlated the spatial distribution of lamin A with 3D nuclear envelope deformation, revealing that local areas of the nuclear envelope with higher expression levels of lamin A have correspondingly lower local deformations, and that increased lamin A expression levels result in a narrower distribution of smaller deformations. These findings describe the complex dispersion of nuclear deformations as a function of lamin A expression and distribution and implicate a role in mechanotransduction.

## Introduction

The nucleus is one of the most crucial organelles and the storehouse of DNA in the cell, integrating diverse biochemical cues for proper cell function^1^. In addition to biochemical cues, the nucleus has an increasingly clear role as a mechanosensory structure in the cell^2^. Nuclear mechanotransduction is a process by which the nucleus detects mechanical cues and converts them to biochemical signals, which alters cell physiology. As a mechanosensor, the nucleus’ stiffness determines specific deformations in response to applied stresses. The nucleus is notably stiffer than the cytoplasm (0.5-3kPa)^3^, with an effective Young’s modulus ranging from 1–10kPa^4,5^; while these values appear relatively consistent in a given cell line^6^, multiple factors may cause nuclear stiffness to change significantly. For example, the stiffness of nuclei varies 5-fold during cell division^7^, in stem cells the stiffness can increase 6-fold over the differentiation process^8^, and in many types of cancers the structure of the nucleus is altered and the stiffness is vastly reduced^9^. Many of these morphological and mechanical changes are the result of the changes in nuclear envelope architecture^10^.

The nuclear envelope is composed of double-membraned bilipid layers with a 30–50nm perinuclear space between the membranes. The outer nuclear membrane is connected to the endoplasmic reticulum, whereas the inner membrane is held in place by a consortium of more than 50 membrane proteins. Some of the key proteins present in the inner nuclear membrane include emerin, SUN1, SUN2, lamins, and lamin associated proteins^11^. Lamins are type V intermediate filaments localized under the nuclear membrane. The inner membrane is held in place by the underlying nuclear lamina, which is mainly composed of A type and B type lamins. Lamin A has a profound effect on the stiffness of the nucleus, but lamin B shows very little or no effect on the nuclear stiffness, thus making lamin A a dominant component in nuclear mechanics^12^.

Numerous studies have illustrated the significance of lamin A in determining the stiffness of the nucleus. Suppression or knockdown of lamin A increases nuclear compliance, and nuclei in lamin A knockout cells display 30% to 50% more deformation than WT when probed with substrate stretch or magnetic microrheology, respectively^13^. Previous studies have also shown that nuclei in lamin A knockout cells are more fragile, with a greater susceptibility to rupture under increased intranuclear pressure^14^. Nuclei isolated from cells transfected with shRNA for lamin A showed 1.5–1.7 times more bead displacement compared to wild type nuclei in magnetic bead microrheology, indicating the role of lamin A in determining the stiffness of the nucleus^15^. Conversely, over-expression of lamin A stiffens nuclei: when 3T3 fibroblasts were transfected with lamin A and seeded on vertical nanopillars, showed 40–50% less nanopillar induced deformation in both intact and isolated nuclei^14^. Lamin A expression and nuclear stiffness also directly affect the behavior of the cell; as an example, lamin A overexpression hinders 3D cell migration through micropores^16^; however, it seems to facilitate 2D migration as wild type cells are marginally faster than lamin A knockout cells^17^. While lamin A expression clearly influences nuclear mechanics and cell behavior, the vast majority of these studies have looked at qualitative treatments of lamin A, i.e., parsing cells into groups of overexpression, wild type, or knock-down. However, the quantitative scaling of nuclear stiffness and corresponding nuclear deformation as a function of lamin A expression remains unknown.

The expression of lamin A is highly variable, and changes in different stages of cell cycle^18^, in stem cell differentation^19^, and as a function of microenvironment stiffness^20^. In addition to overall expression levels, the distribution of lamin A appears heterogeneous with the formation of foci and honeycomb patterns observed in the nuclear envelope^21^. It remains unknown how the spatial heterogeneity of lamin A affects nuclear deformation. Studies to date have focused on comparing gross nuclear stiffness changes in ensemble population lamin A expression levels via over-expression or suppression^22^, but have not examined the quantitative dependence of nuclear stiffness on lamin A expression, or the impact of the spatial distribution of lamin A of deformation. To understand the quantitative impact of lamin A structure on nuclear mechanics, here we measure the deformability and bulk moduli of nuclei as a function of lamin A expression. We identify a mechanical anisotropy of bulk compression, leading to increasing relative compliance in the major, minor, and z axes, respectively. At the sub-micron level, we quantify the spatial distribution of lamin A and show that dense lamin A structures attenuate local deformation, providing heterogeneous nuclear deformations even under uniform nuclear stresses.

## Methodology

### Quantification of lamin A expression using antibodies and lamin A chromobody

To quantify the total lamin A expression in the nucleus, the NIH 3T3 fibroblast cells were stably transfected with GFP tagged lamin A chromobody^23^ (Chromtek). To testify if the chromobody is an accurate quantitative metric of lamin A expression and distribution, we compared its fluorescence with antibody staining (Atto-647N Sigma) as a gold standard (see supplementary information Fig. S1). For this, lamin A chromobody transfected cells were seeded onto gridded coverslips, fixed, permeabilized and stained with Atto-647 lamin A antibodies and then imaged with a 63X/1.4 NA oil immersion objective on a confocal microscope (Leica SP8). Fluorescence colocalization was determined using Pearson’s correlation coefficient of 0.88±0.04 for 10 cells (see supplementary information Fig. S2).

### Cell culture & modulating nuclear lamin A expression

NIH 3T3 fibroblast cells were cultured following standard protocols in DMEM media supplemented with 10% FBS and 1% Penicillin-Streptomycin antibiotic. All cells were stably transfected with GFP tagged lamin A chromobody, which labels the total lamin A present on the nuclear membrane without affecting its expression. To over-express lamin A in intact cells, we transfected cells with m-Cherry tagged plasmid DNA for lamin A which was a gift from Michael Davison (Addgene plasmid # 55068), whereas to suppress lamin A expression we transfected the cells with RFP tagged inducible shRNA construct for lamin A (Dharmacon).

### Quantification of nuclear bulk moduli

The cells were synchronized in 0.1% FBS containing media for 18 hours to eliminate the contribution of cell cycle on lamin A expression and distribution. They were then attached on coverslips and exposed to media with 1%, 2.5%, 5%, 7.5% and 10% (wt/wt) 400 Da polyethylene glycol (PEG 400, Sigma) thus, exerting 847.7kPa, 936.5kPa, 1099.2kPa, 1290.3kPa, and 1514kPa osmotic stresses respectively for 25 minutes to reach an equilibrium of compression^24^ (see supplementary information Fig. S3), followed by acquiring confocal microscopy XYZ-stacks (63x 1.4NA, Leica SP8). The cells compress due to the osmotic pressure exerted by the hypertonic solution causing an efflux of water^25^, and this change in volume is measured and used to determine the bulk modulus by the relation B = -ΔP/ (ΔV/V) where, B= bulk modulus, ΔP = osmotic pressure, ΔV= change in volume and V = original volume^6^.

### 3-D volume measurement of nuclei

XYZ stacks of GFP tagged lamin A chromobody cells were imaged using a 63X/1.4 NA oil immersion objective on a Leica SP8 confocal microscope with a z-step size of 0.3µm. 3-D visualization and measurement were carried out using ImageJ software. Before 3-D measurement, the nuclear images were deconvolved using the ‘Iterative deconvolve 3D’ plugin in ImageJ. We measured nuclear volumes by counting the number of voxels of the thresholded nuclei and then multiplied it by the size of each voxel to get the volume of the thresholded region. To measure the major, minor and z-axis deformations, we acquired XYZ stacks for different time intervals post-PEG addition. The stacks were summed in the Z dimension using ‘z-project’ plugin in image J, followed by fitting the z-projections into an ellipse and calculating the major and minor axis for different time intervals.

### Nuclear strain mapping

To measure the local strain on the nuclear membrane, we captured images of the GFP tagged lamin A chromobody nucleus pre and post-exposure to 2.5% PEG 400 (936.5kPa). To quantify the local deformation, a custom-made MATLAB code^26^ was slightly modified and used. Briefly, the code utilizes a fast Fourier transform (FFT) based cross-correlation formulation in conjunction with the iterative deformation method (IDM). FIDVC calculates the displacement between the images of GFP tagged nucleus by tracking the pixel intensities of chromobody fluorescence. We mapped these local displacements with the lamin A expressions in 3-D space to determine the relationship between lamin A spatial distribution and nuclear deformation.

## Results

### Lamin A stiffens the nucleus

To measure the bulk compressibility of the nuclei, we exposed cell nuclei of varying lamin A expression with different osmotic pressures using 400 Da PEG. Since PEG is impermeable across the cell membrane, it creates a hyperosmotic environment for the cell and nucleus resulting in their compression (Fig. 1a). This compression is due to the expulsion of water as the cell and nuclei regain their original volume when returned to isotonic media^6,25^. Previous studies have shown that under external osmotic pressure nuclear volume changes along with the concentration of intracellular macromolecules, thus exerting compressive forces on it^6,25^. The pressure-volume curve for nuclei of wild type NIH 3T3 fibroblast cells at different PEG 400 concentrations exerting different osmotic pressures shows that the nuclear compression was higher in low lamin A expressing cells and vice-versa (Fig. 1b)

**Figure 1.**
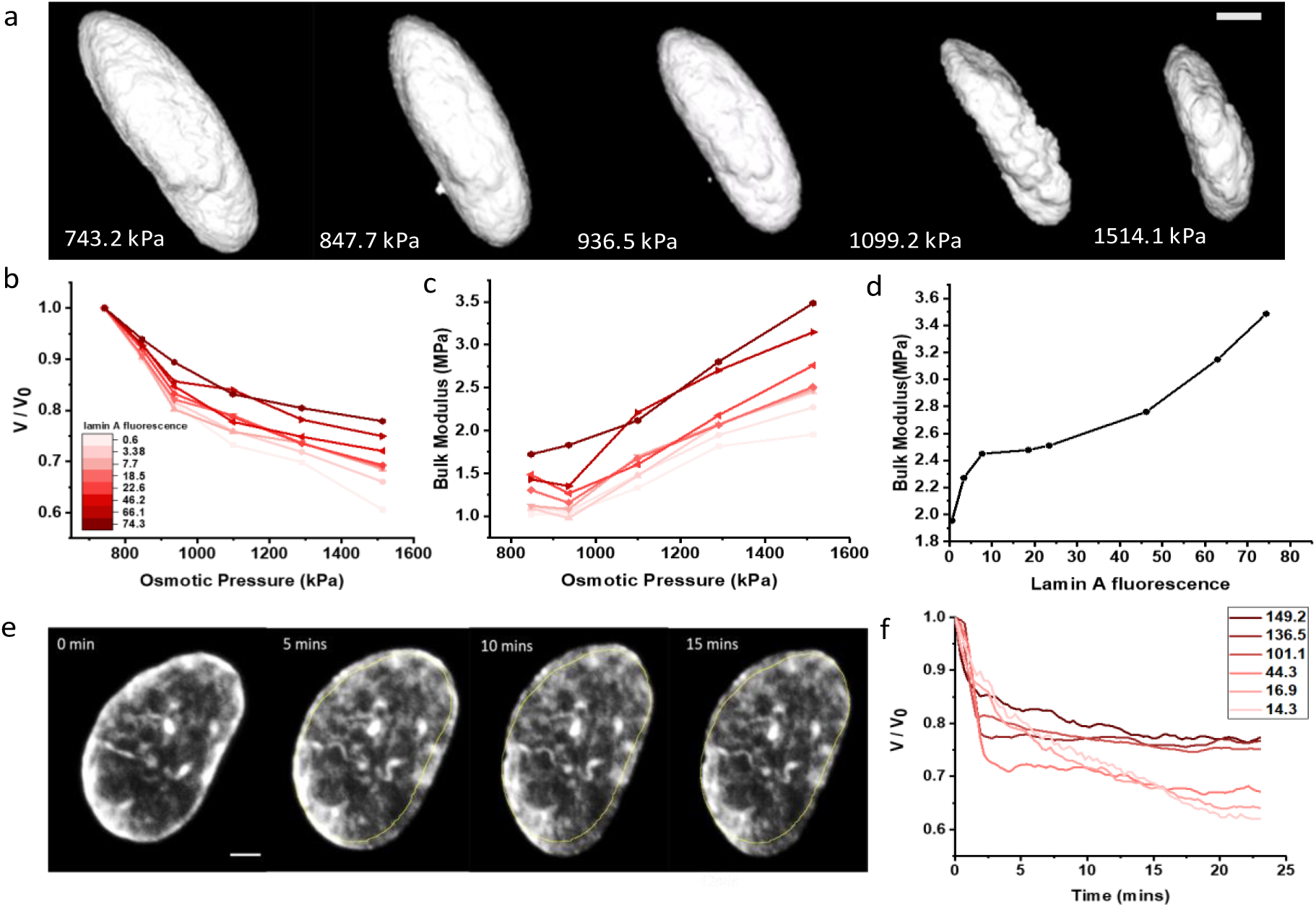
Hyperosmotic compression of the nucleus in 3T3 Fibroblast cells. **a**, 3D reconstruction of an example nucleus within a cell at osmotic pressures of 743.2 and 1514.1kPa (scalebar=3µm). **b**, Normalized nuclear volumes of 3T3 cells transfected with lamin A chromobody for wild type cells showing that stiffness increases as the protein expression increases (fluorescence values in arbitrary units). **c**, Bulk moduli of 3T3 cells transfected with lamin A chromobody for wild type showing that stiffness increases as the protein expression increases (n=8). **d**, Nuclear stiffnes in terms of bulk moduli increasing as a function of lamin A fluorescence. **e**, Example 3T3 fibroblast nuclei showing change in nuclear volume with time after PEG addition (scalebar = 2μm). **f**, Change in nuclear volume as a function of time under 1514.1kPa osmotic pressure as a function of lamin A fluorescence.

This is also reflected in the bulk moduli calculated from the change in the nuclear volume where the cells expressing more lamin A are relatively stiffer. The bulk modulus for the highest lamin A expressing cell at the highest exerted osmotic pressure is 3.48Mpa (Fig. 1c). A linear relationship could be seen when we plotted bulk moduli as a function of lamin A fluorescence clearly a direct relationship between lamin A expression and nuclear rigidity (Fig. 1d). We also observed the nuclear volume as a function of time for different lamin A expression cells after PEG addition and show that below lamin A fluorescence of 16.9 the rate of nuclear compression was more uniform, whereas in nuclei with lamin A expression more than 16.9 showed a sudden fall in nuclear volume followed by a very slow compression indicating different mechanical behavior of the nuclei at below a certain level of lamin A expression (Fig. 1e & 1f).

### Nuclear stiffness is anisotropic

To measure the anisotropic nuclear deformation, we exposed the cells transfected with lamin A chromobody to PEG and imaged with confocal microscopy (Leica SP8) after 25 minutes of PEG addition. Our results reveal strong anisotropic nuclear deformation where the deformation in the z-axis was highest, followed by minor and major axis (Fig 2a). To examine the dynamics of nuclear deformation under stress, we imaged the nuclei every 60 seconds after PEG addition. Here, the nucleus quickly flattens in the z-axis, resulting in a transient expansion along the major and minor axes, which starts to contract after 5 minutes (Fig. 2b). This shows a time dependency to nuclear compression after PEG addition. Also, the deformation in z-axis was inversely related with lamin A expression but the deformation along major and minor axis did not show any relationship with lamin A expression (Fig. 2c).

**Figure 2.**
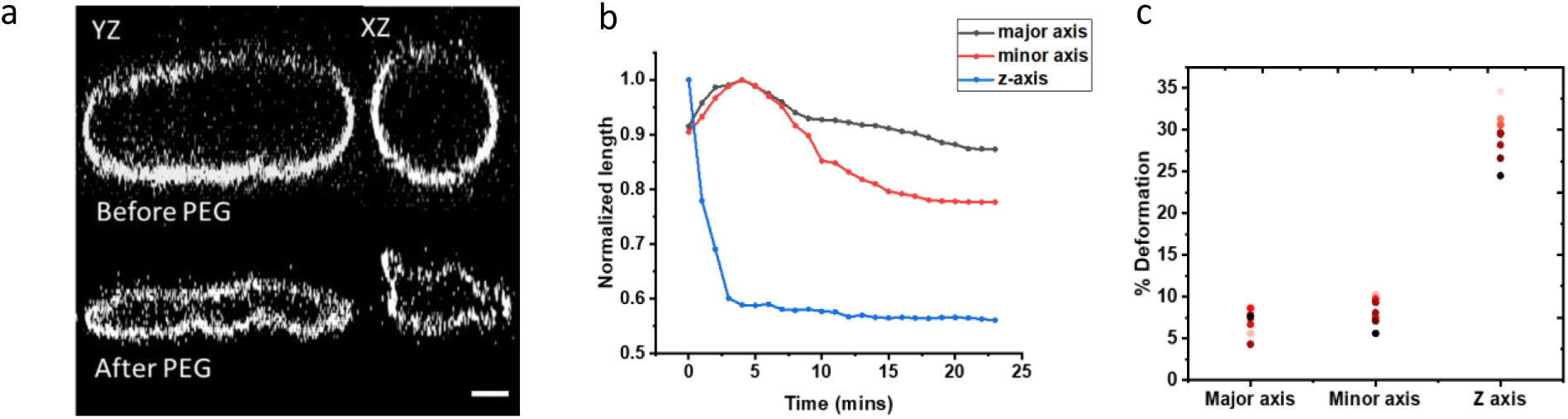
Anisotropy in nuclear compression under PEG. **a**, Example of a wild type nuclei in Y-Z and X-Z plane before PEG addition (top) and after PEG addition (bottom) showing large deformation in the z-axis (scalebar = 3µm). **b**, Example of a 3T3 fibroblast nucleus showing that the z-axis gets compressed instantly after PEG addition whereas major and minor axes expand in the beginning followed by reduction in the length showing anisotropy in nuclear compression with time. **c**, Nuclear deformation percentage along the major, minor and z-axis in wild type nuclei for different lamin A expression (n=7) showing higher deformation along z-axis compared to the major and minor axis.

### Lamin A density is spatially heterogeneous and leads to a variation in local compliance

While overall lamin A expression levels are typically considered for overall nuclear compliance, we find that the spatial distribution of lamin A is not uniform in the nuclear envelope (Fig. 3a and 3b). This inhomogeneity also increases with overall lamin A expression, as revealed by plotting the variance of local lamin A fluorescence intensity as a function of overall expression (Fig. 3c, supplementary information Fig. S5). The histogram of local lamin A fluorescence shows narrow and high peak at lower lamin A fluorescence levels but the peak broadens and flattens with increase in local lamin A fluorescence levels indicating that lamin A distribution gets wider or heterogenous with increase in lamin A expression (Fig. 3d). We also observed the bin span from Fig. 3d as a function of total lamin A expression to show that the range of lamin A fluorescence covered increased with lamin A expression showing a greater heterogeneity and vice-versa (Fig. 3e).

**Figure 3.**
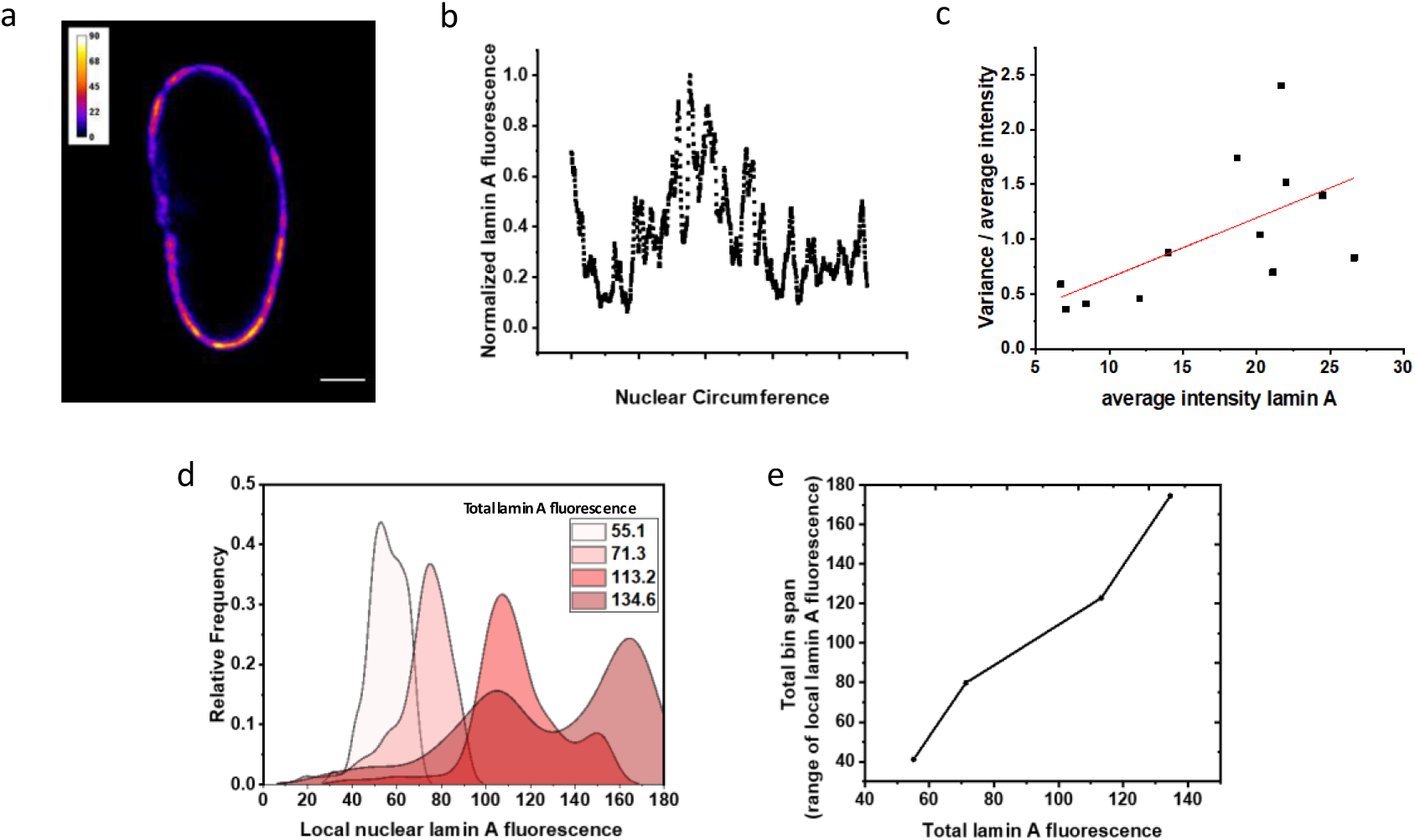
Lamin A is spatially heterogeneous in the nucleus. **a**, Example 3T3 fibroblast cells transfected with lamin A chromobody showing variability in the spatial distribution along a cross-section nuclear membrane (scalebar = 3µm). **b**, Quantification of lamin A heterogeneity along the nuclear membrane circumference in panel **a. c**, Normalized variance of lamin A chromobody fluorescence as a function of total lamin A expression in different nuclei shows that higher expression of lamin A increases the heterogeneity of distribution (n=12). **d**, The histogram gets wider with increasing total lamin A fluorescence showing a wider range of local lamin A fluorescence or heterogeneity when overall lamin A expression increases. **e**. The bin span correlates linearly with lamin A expression showing a more heterogeneous and wider distribution at high lamin A fluorescence and vice-versa.

We then characterized the relationship between local lamin A expression and local nuclear deformation by quantifying the lamin A distribution via fluorescence and the nuclear deformation map in the X-Y and Z plane along the nuclear membrane using FIDVC (Fig. 4a, 4b, and 4c). These images suggest an inverse relationship between spatial variations of lamin A and deformation. To quantify the spatial variations, we plotted lamin A fluorescence with respective deformations along the nuclear circumference, showing the deformation as a function of the fraction of lamin A present at the nuclear membrane (Fig. 4d and Fig. 4e). We observed the least nuclear deformation in regions of highest lamin A expression and maximum deformation at regions of lowest lamin A expression. The ratio of applied osmotic stress to local nuclear deformation as a function of the lamin A expression yields the effective contribution of lamin A to nuclear stiffness; this reveals that the stiffness increases as lamin A expression increases locally (Fig. 4f). While this finding is consistent in all cells measured, some cells also presented low deformations in regions of low lamin A expression (see supplementary information Fig. S6), which may be attributed to the mechanical anisotropy of the underlying chromatin^27^. This may also contribute to the observed mechanical anisotropy of the nucleus along the major and minor axes of the nucleus, where the minor axis is deformed slightly more than the major axis (Fig. 4g). The correlation coefficient between local deformation and local lamin A fluorescence for 10 cells was found to be negative showing a moderate anti-correlation between them but we did not find any relation of the correlation coefficient values with the total lamin A expression (Fig. 4h). This variation in anticorrelation could be due to the effect on the underlying chromatin.

**Figure 4.**
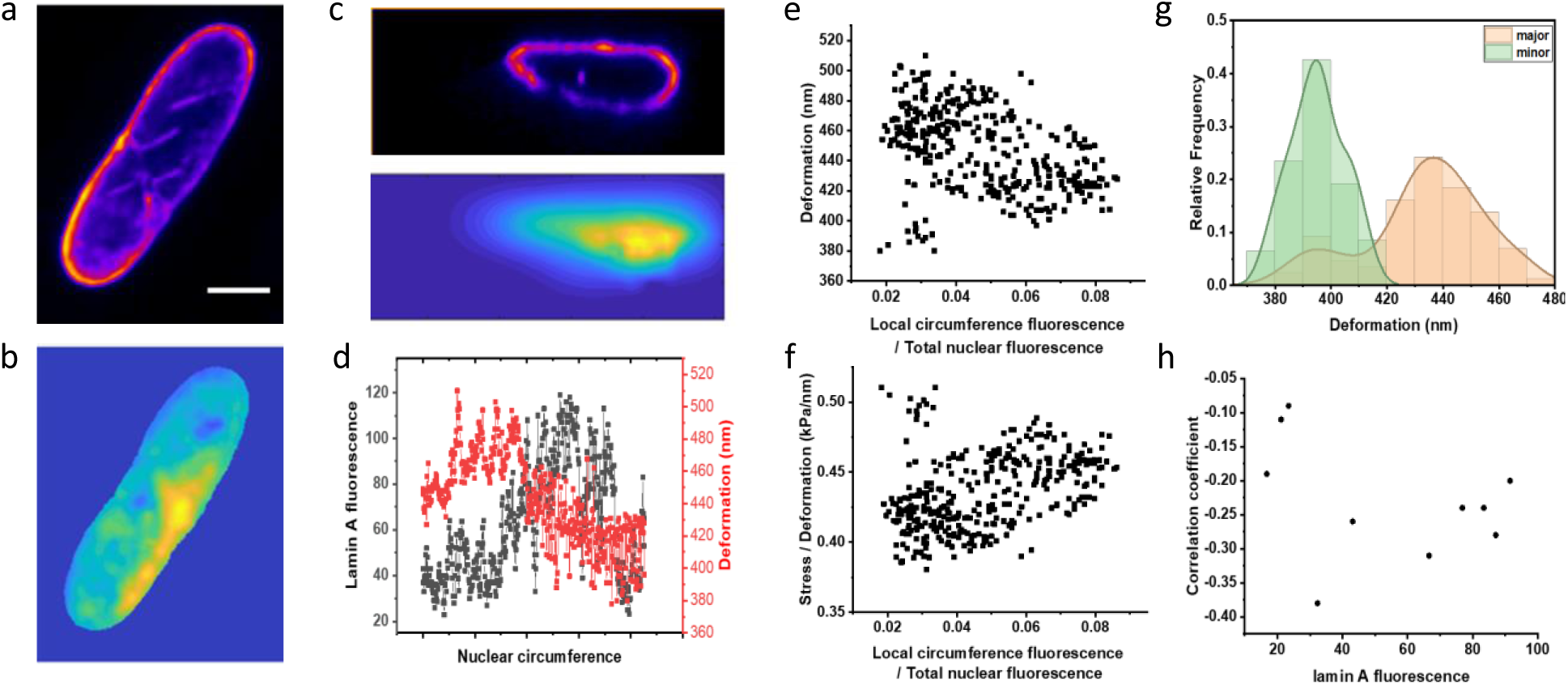
Local nuclear deformation as a function of lamin A distribution in 3T3 fibroblasts. **a**, Lamin A distribution along the nuclear membrane in X-Y plane. **b**, Example strain X-Y plane map of a 3T3 nucleus under 936.5kPa stress (scalebar = 5µm). **c**, Lamin A distribution (top) and Y-Z strain map (bottom) along the nuclear membrane. **d**, Lamin A fluorescence and nuclear deformation along the nuclear membrane. **e**, Deformation plotted against the ratio of local lamin A fluorescence at the membrane and total nuclear fluorescence. **f**, Ratio of stress to deformation (effective stiffness) plotted against the ratio of local lamin A fluorescence at the membrane and total nuclear fluorescence. **g**, Statistical distribution of overlaid major and minor axis. **h**, The correlation coefficient between local deformation and local lamin A fluorescence was negative showing anticorrelation but with no relationship with total lamin A expression.

## Discussion

Here, we have quantitatively examined the role of lamin A in nuclear mechanics. Using osmotic pressure, we applied precisely controlled stresses and measured global volumetric changes. We find that the overall bulk modulus of the nucleus increasing with expression levels of Lamin A. Decomposing nuclear compression, we identify anisotropic deformation of the nucleus showing maximum deformation in z-axis followed by minor axis and then major axis, respectively. Inspecting the distribution of lamin A more closely, we found distinct spatial variations in concentration within the nuclear envelope; interestingly, the heterogeneity of lamin A density increased with overall expression level. We suspected that these spatial variations in concentration would lead to similar spatial variations in nuclear stiffness. By quantifying the spatial variations of lamin A density and comparing this with a sub-micron resolution deformation map of the nucleus, we indeed found a strong inverse correlation between local sub-micron lamin A concentration and nuclear deformation, demonstrating that the spatial variation of lamin leads to heterogeneous nuclear membrane deformation. Also, the local deformations along the minor axes of the nuclei were found to be larger than the corresponding major axes, which was consistent with our previous observation, as seen in the bulk deformation of the nuclei.

The expression and distribution of lamin A are critical in nuclear mechanics, with variations or abnormalities in lamin A affecting nuclear stiffness leading to altered nuclear deformation. Pathological conditions such as HGPS^28^ and EDMD^29^ are related to a mutation in the *lmna* gene suggesting a potential role of nuclear deformation in nuclear mechanotransduction in the cell, but the relationship between nuclear deformation and how it changes the gene expression is not clear. The strongest evidence to this hypothesis comes from a recent study, where it is shown that when the nucleus is physically compressed, there is translocation of an enzyme Histone deacetylase 3 (HDAC3) inside the nucleus. The role of this enzyme is to remove the acetyl group present on histone residues which neutralize the positive charges present on the histones and inhibits its interaction with the negatively charged DNA resulting in an open conformation of chromatin known as euchromatin (non-compact expressing chromatin conformation). Thus, when the nucleus is mechanically compressed, HDAC3 translocates in the nucleus to remove the acetyl group on the histone leading to a tighter histone-DNA interaction promoting the non-expressing heterochromatin (highly compact non-expressing chromatin) conformation of the chromatin^30^.

Another factor contributing to nuclear stiffness is the tethering of chromatin itself with the nuclear lamina^31^. Since the chromatin can be present as heterochromatin and euchromatin, it contributes to the mechanical anisotropy of the nuclear envelope^32^. While lamin A dominates nuclear stiffness under high mechanical stress, under small stresses, chromatin has a crucial role as well^27^. More condensed heterochromatin regions are stiffer and experience less deformation compared to euchromatin^33^, making chromatin dynamics additionally important in nuclear mechanotransduction. Nevertheless, the physical association between the nuclear lamina and the chromatin, as well as the relationship between nuclear deformation and its direct effect on epigenetic changes are still unclear and could prove to be essential in relating nuclear mechanics with various pathological conditions.

Previous mechanotransduction studies have shown that mechanical forces can lead to translocation of mechanosensitive proteins such as YAP from the cytoplasm to the nucleus under nuclear tension^34^, which affects properties like cell proliferation and differentiation in stem cells. Lamin A could prove to be a critical factor in elucidating this relationship as it directly affects the nuclear stiffness and its deformation. By manipulating lamin A expression, relating the change in nuclear mechanics, and measuring downstream effects, we may better understand how nuclear stiffness determines cellular fates, thus providing novel mechanics-centered strategies to correct defects in diseases.

## Acknowledgments

The authors thank members of Christian Frank’s lab, University of Wisconsin, for assistance with MATLAB FIDVC. AJE acknowledges support from NSERC (RGPIN/05843-2014, EQPEQ/472339-2015, RTI/00348-2018), CIHR (Grant # 143327), and the Canadian Foundation for Innovation (Project #32749).

