## Supplementary Information for "Spatial distribution of lamin A determines nuclear stiffness and stress-mediated deformation"

**Quantification of lamin A expression using antibodies and lamin A chromobody:**

To visualize the total lamin A present inside the cells, they were fixed and stained with Atto-647N tagged lamin A antibodies. Lamin A chromobody is a substitute for antibodies in live cells as it tags the whole lamin A without affecting the lamin A expression or cellular physiology. Fig. S1 shows the similarity in staining by lamin A antibodies and lamin A chromobody.

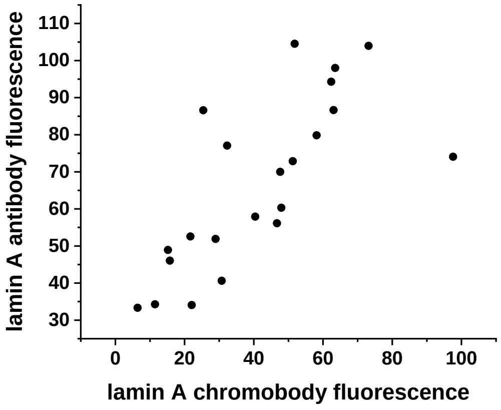


**Figure S1**: Relationship between overall antibody and chromobody fluorescence across unique cells(n=18)

Besides, the total fluorescence, results show that the relationship was linear even across the membrane of the same nucleus. To establish a more quantitative correlation, colocalization analysis was performed. Fig. S2a shows images of lamin A antibody (red) and lamin A chromobody (green). Fig. S2b and S2c show that the correlation exists even across the nuclear membrane quantifying lamin A heterogeneity. Fig. S2d shows an example of the colocalization analysis between lamin A chromobody and lamin A antibody. The colocalization analysis was done for 10 cells with an average Pearson's R-value of 0.88±0.04 which indicates that both the fluorophores are strictly colocalized.

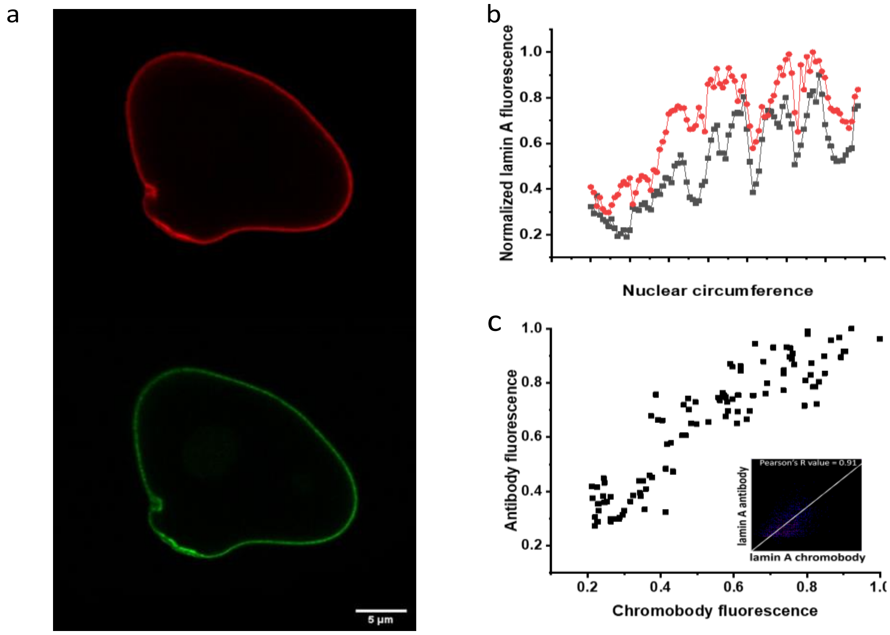


**Figure S2**: **Spatial correlation between lamin A antibody and lamin A chromobody in the cell. a,** Nucleus stained with antibody tagged with Atto-647N (top) and GFP-tagged lamin A chromobody (bottom) (scalebar = 5µm). **b,** lamin A antibody (black) and lamin A chromobody (red) fluorescence of the same points along the nuclear circumference. **c,** Correlation between antibody and chromobody fluorescence of the same points along the nuclear circumference and an example of colocalization analysis of lamin A chromobody and lamin A antibody has been shown in the inset.

**Hyperosmotic nuclear compression:** 3d reconstruct of a nucleus under PEG 1%, 2.5%, 5%, and 10% 400 Da polyethylene glycol (PEG 400, Sigma) thus, exerting 743.2kPa, 847.7kPa, 936.5kPa, 1099.2kPa, and 1514kPa osmotic stresses respectively for 25 minutes has been shown in Fig. S3.

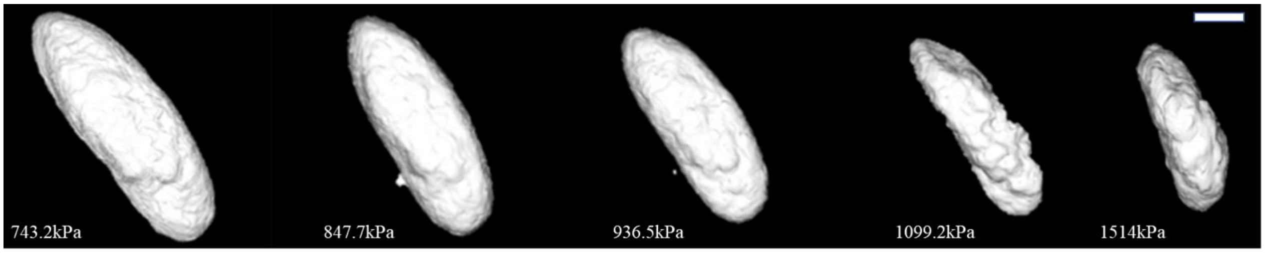


**Figure S3:** 3D Nuclear deformation of a nucleus under 743.2kPa, 847.7kPa, 936.5kPa, 1099.2kPa, and 1514kPa osmotic stresses (scale bar = 3µm).

**Anisotropy in nuclear compression:** The nuclear deformation of 15 cells along the major, minor and z-axis has been shown in Fig. S4a. Fig. S4b shows z projections of z-stacks at different time intervals (0 minutes, 3 minutes, 6 minutes, 9 minutes, 12 minutes, 15 minutes, 18 minutes) after the addition of PEG. There is a sudden reduction in z-axis with simultaneous stretching in the major and minor axis which starts to reduce after 6 minutes of PEG addition clearly showing the anisotropic behavior in nuclear compression after PEG addition.

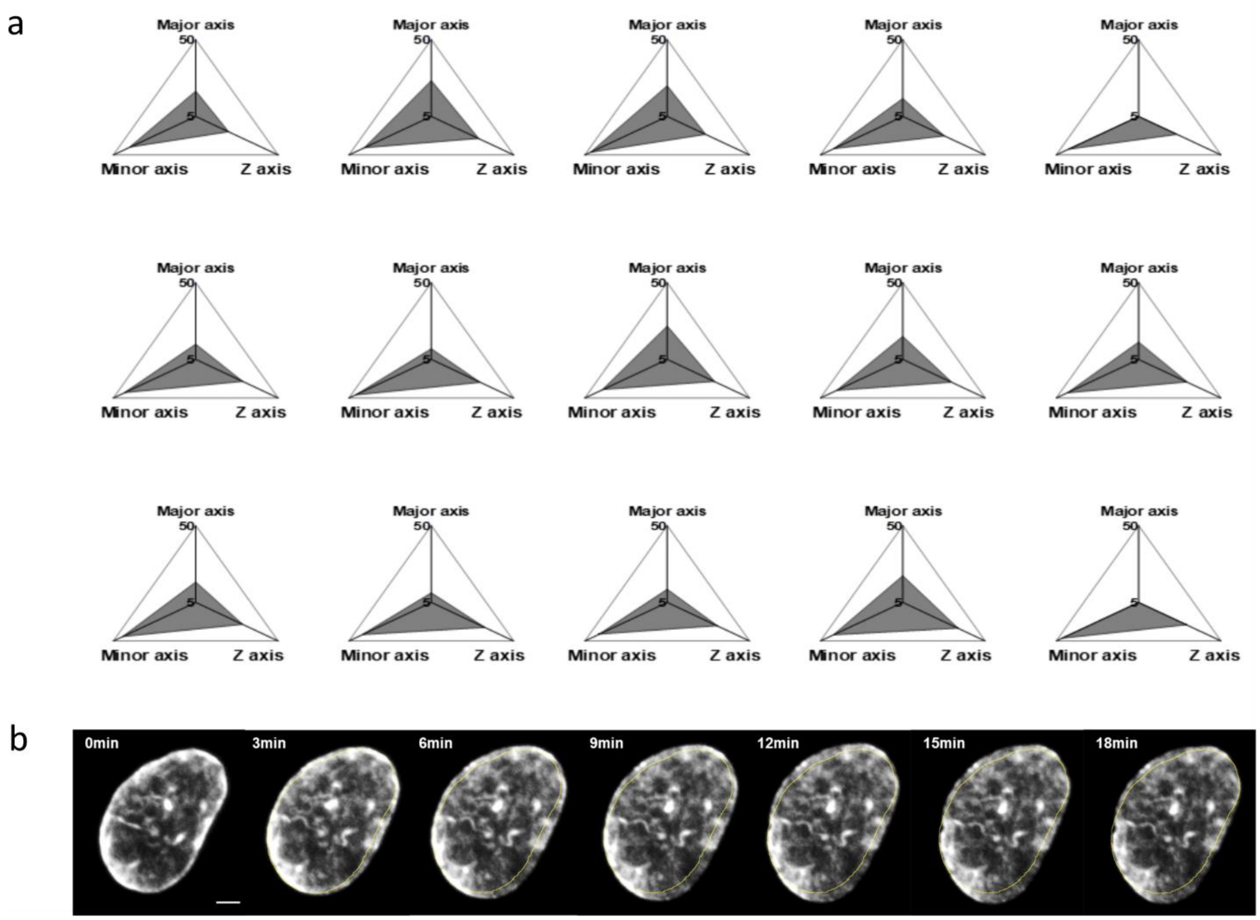


**Figure S4: Anisotropy in nuclear compression. a,** Nuclear deformation of 15 nuclei along major, minor and z-axis showing the largest deformation along the z-axis. **b,** Z-projections of wild type nucleus stained with lamin A chromobody at different time intervals with the original zero pressure image overlaid as a visual aid (scalebar = 2μm).

**Lamin A spatial heterogeneity:** Images of nuclei transfected with lamin A chromobody are shown in Fig. S5a. As the lamin A fluorescence increases, the heterogeneity can be seen to increase as well. To eliminate the role of intensity artifacts skewing this relation, the fluorescence of images was multiplied by 1.5x and 2x, and the normalized variance thus calculated was plotted as a function average intensity respectively as shown in Fig. S5b. The slope was found to be 0.05 and did not change with artificial fluorescence increase, demonstrating measured increases in lamin A heterogeneity are due to changes in overall expression.

a

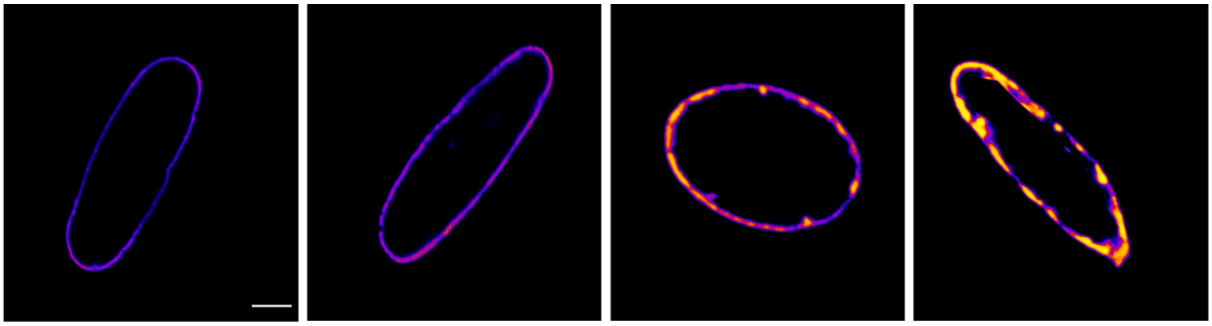

b

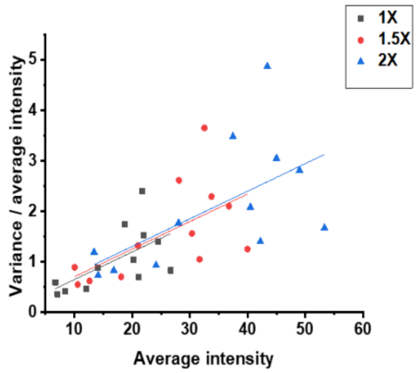


**Figure S5: Local nuclear deformation as a function of lamin A protein in 3T3 fibroblast cells. a,** Increasing lamin A heterogeneity with an increase in lamin A fluorescence (scale bar = 3μm)**. b,** Normalized variance of lamin A chromobody fluorescence (n=12) showing that spatial heterogeneity of lamin A increases with expression levels and the slope does not change after multiplying by factors of 1.5 and 2 times showing no role gain in lamin A heterogeneity.

### Nuclear strain mapping

Fig. S6a and S6b show additional examples of nuclear deformation and lamin A distribution. Fig. S6c shows three different populations. The first one being the nuclear deformation maximum at the points where lamin A expression is minimum. The second showed regions with a high level of lamin A showing reduced deformation and in the third population there are some regions showing low deformation at some points with intermediary levels of lamin A. The first two are expected but the third population might be a result of the mechanical anisotropy due to the underlying chromatin.

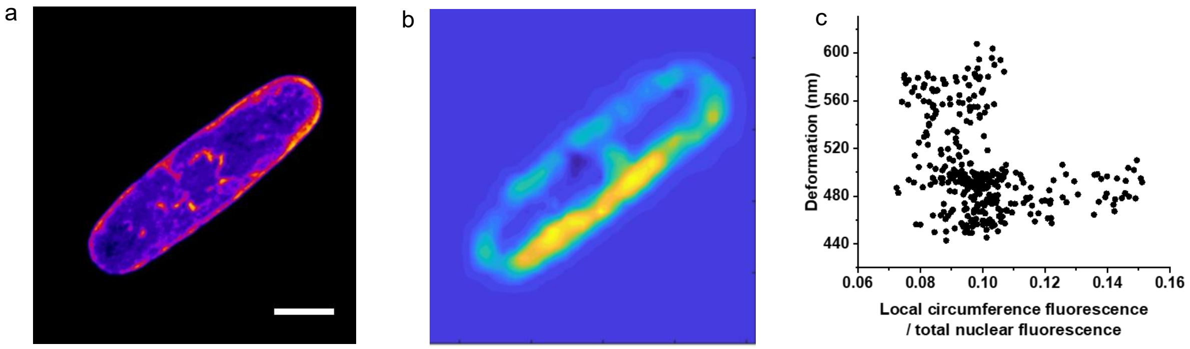


**Figure S6: Local nuclear deformation as a function of lamin A protein in 3T3 fibroblast cells. a,** lamin A distribution along the nuclear circumference in the X-Y plane (scale bar = 5μm). **b,** Nucleus strain map in the X-Y plane. **c,** Nuclear deformation as a function of lamin A fluorescence.
